# A simple theory for finding related sequences by adding probabilities of alternative alignments

**DOI:** 10.1101/2023.09.26.559458

**Authors:** Martin C. Frith

**Affiliations:** Artificial Intelligence Research Center, AIST, Tokyo, Japan; Department of Computational Biology and Medical Sciences, University of Tokyo, Chiba, Japan; Computational Bio Big Data Open Innovation Laboratory, AIST, Tokyo, Japan

## Abstract

The main way of analyzing genetic sequences is by finding sequence regions that are related to each other. There are many methods to do that, usually based on this idea: find an alignment of two sequence regions, which would be unlikely to exist between unrelated sequences. Unfortunately, it is hard to tell if an alignment is likely to exist by chance. Also, the precise alignment of related regions is uncertain. One alignment does not hold all evidence that they are related. We should consider alternative alignments too. This is rarely done, because we lack a simple and fast method that fits easily into practical sequence-search software. Here is described a simplest-possible change to standard sequence alignment, which sums probabilities of alternative alignments. Remarkably, this makes it easier to tell if a similarity is likely to occur by chance. This approach is better than standard alignment at finding distant relationships, at least in a few tests. It can be used in practical sequence-search software, with minimal increase in implementation difficulty or run time. It generalizes to different kinds of alignment, e.g. DNA-versus-protein with frameshifts. Thus, it can widely contribute to finding subtle relationships between sequences.

## Introduction

Many methods have been developed for finding related parts of nucleotide or protein sequences. They usually follow a standard approach: define positive and negative scores for monomer matches, mismatches, and gaps, then seek high-scoring alignments. Why is this bizarre, intricate approach a good way to find related sequences? Its result depends utterly on the score parameters, so how should we choose them?

The answer to both questions is that the approach is equivalent to probability-based alignment [1, 2]. This uses probabilities of matches, mismatches, and gaps, e.g. their average rates in the kinds of related sequences we wish to find, and seeks high-probability alignments. An alignment’s score is a log probability ratio [3]:

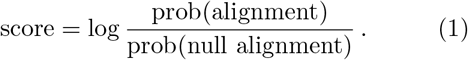

A “null alignment” is a length-zero alignment between the two sequences. A precise definition of prob(null alignment) was given previously [3], and is repeated in this paper’s appendix. A high alignment score indicates high probability with these match/mismatch/gap rates. The base of the log (e.g. log_2_, log_10_) is unspecified, because any base is equally correct. A change of base merely rescales the scores. (In the precursor to this paper [3], log *x* was equivalently written as *t* ln *x*, where *t* is an arbitrary positive constant.) This approach is termed “pairwise local alignment”, where “local” means that the alignment is not necessarily of the whole sequences.

A more direct way to judge whether two sequence regions are related is to calculate their probability without fixing an alignment. In other words, use a score like this:

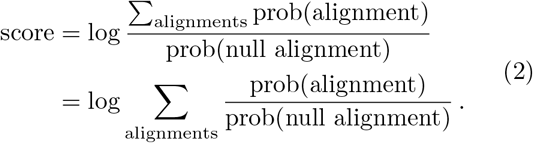

If the sum includes all possible ways of aligning the sequence regions, it gives their total probability with the assumed rates of matches, mismatches, and gaps. Thus, it includes all evidence that the sequence regions are related, while one alignment includes just some evidence.

A similar sum-over-alignments method has been used with great success in the HMMER software [4, 5]. So it is curious that many sequence-search tools developed since HMMER do not use this approach. No-one has described a sum-over-alignments method that fits easily into typical sequence-search software, with minimal increase in implementation difficulty or computational cost. HMMER is too complicated and specialized, e.g. it has an enormous “implicit probabilistic model”, and sequence-length–dependent parameters [4]. HMMER is designed to search a smallish “query” sequence against a large “database”, so it can’t find related parts of two large sequences, e.g. two genomes.

It is important to know whether a similarity score (eqs. 1, 2) is likely to occur by chance between unrelated sequences. The simplest definition is: the probability of an equal or higher score between two sequences of random i.i.d. (independent and identically distributed) monomers. Even this simplest definition, however, is hard to calculate. There is a solution for gapless alignment, in the limit where the sequences are long [6]. Gapped alignment relies on conjecture and simulation. BLAST can calculate it only for a few sets of score parameters, not including realistic DNA scores that distinguish transition (a:g, c:t) from transversion mismatches [7]. Some widely-used aligners do not calculate it at all [8]. The ALP library can calculate it for arbitrary parameters, by complex simulations [9]. Remarkably, HMMER calculates it in an easier way, based on other conjectures. The scope of those conjectures is unclear, e.g. they seem to hold only after tweaking HMMER to use a “uniform entry/exit distribution” [4].

Here is described a minimal modification of standard alignment, which sums probabilities of alternative alignments (eq. 2). It can replace the alignment component of practical sequence-search software, with minimal increase in implementation difficulty or run time. This also makes it easier to calculate probabilities of similarity scores between random sequences, based on a clear conjecture.

### Limitations of previous methods

Many previous studies have summed probabilities of alternative alignments. This section surveys some representatives, to understand why this approach is rarely used to find related parts of sequences.

One line of research considered substitutions, insertions, and deletions occuring over time, so that match/mismatch/gap probabilities depend on the length of time [10–14]. These methods achieve several aims. Given two related sequences, they can: infer substitution/insertion/deletion rates per unit time, find the most likely alignment, and report the probability that any pair of monomers descend from a common ancestor. Furthermore, Hein et al. used a log likelihood ratio to compare two hypotheses, that two whole sequences are related or not:

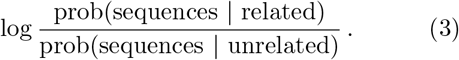

The numerator is a sum of probabilities of all possible alignments between the whole sequences. These studies achieved exact mathematical solutions only for simple models of sequence evolution, which do not fit real sequences well. In particular, it seems hard to allow insertion and deletion events of length *>* 1. To my knowledge, this approach has never been used to find related parts of sequences.

Another line of research (including the present study) simplifies things by ignoring time-dependence, and directly using probabilities of matches, mismatches, and gaps. Allison et al. [15] estimated such probabilities from a pair of related sequences, allowing complex gap-length distributions. They also used a log likelihood ratio to compare hypotheses that two whole sequences are related or not.

In a pioneering study, Bucher & Hofmann used a log likelihood ratio to compare two hypotheses, that two sequences have a related segment or not [16]:

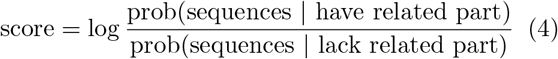

Each hypothesis has probabilities for the lengths of the two sequences. Their two hypotheses are identical in this regard. Thus, the sequence lengths never favor either hypothesis. This is arguably unnatural, biologically and mathematically. Biologically, if related regions are equally likely to occur per unit sequence length, longer sequences are more likely to have related regions. For example, a longer chromosome is more likely to contain a mobile DNA element. Mathematically, they need two summation algorithms. The likelihood ratio becomes a fraction where the numerator is a sum of exponentiated alignment scores, and the denominator is another sum. This denominator increases with increasing sequence lengths [17, fig. 2]. So scores of related regions decrease with increasing lengths of sequences containing them.

**Figure 1:**
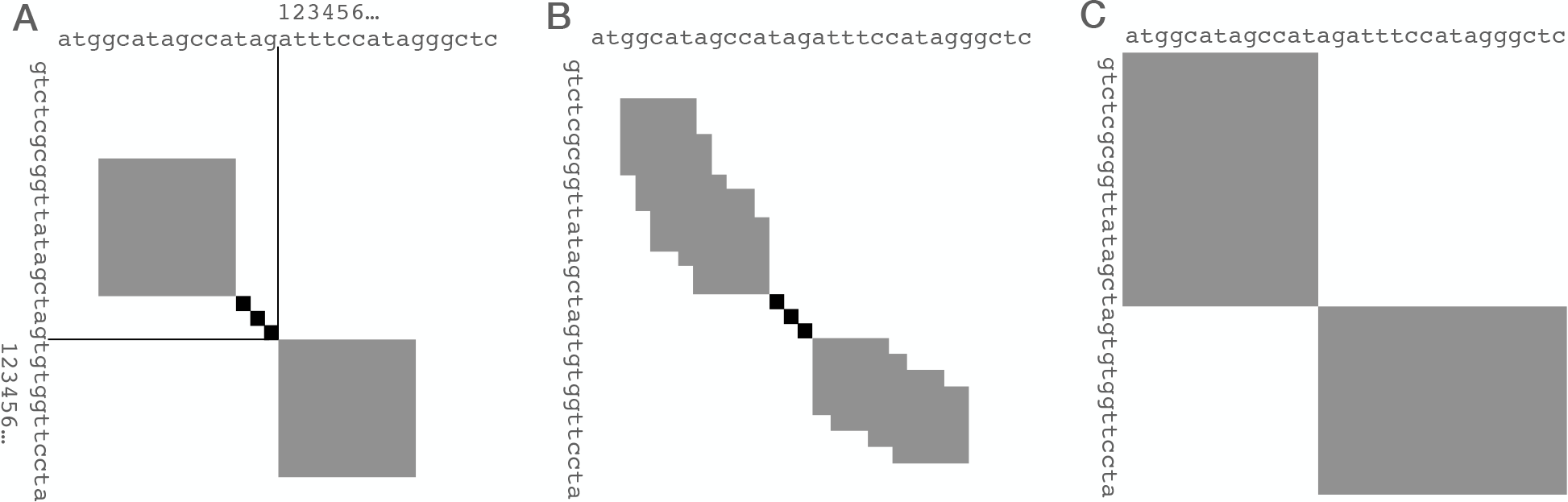
Some ways of extending an alignment in both directions from a small “core” alignment (black). In each direction, an optimum alignment extension is sought from among all possible ones within the gray area. **A** Simple toy method: the gray areas are blocks of predefined size *n* × *n*. **B** Realistic example: the gray areas are adjusted as the calculation proceeds, attempting to follow good alignments. **C** Unlimited extension from zero-size core.

**Figure 2:**
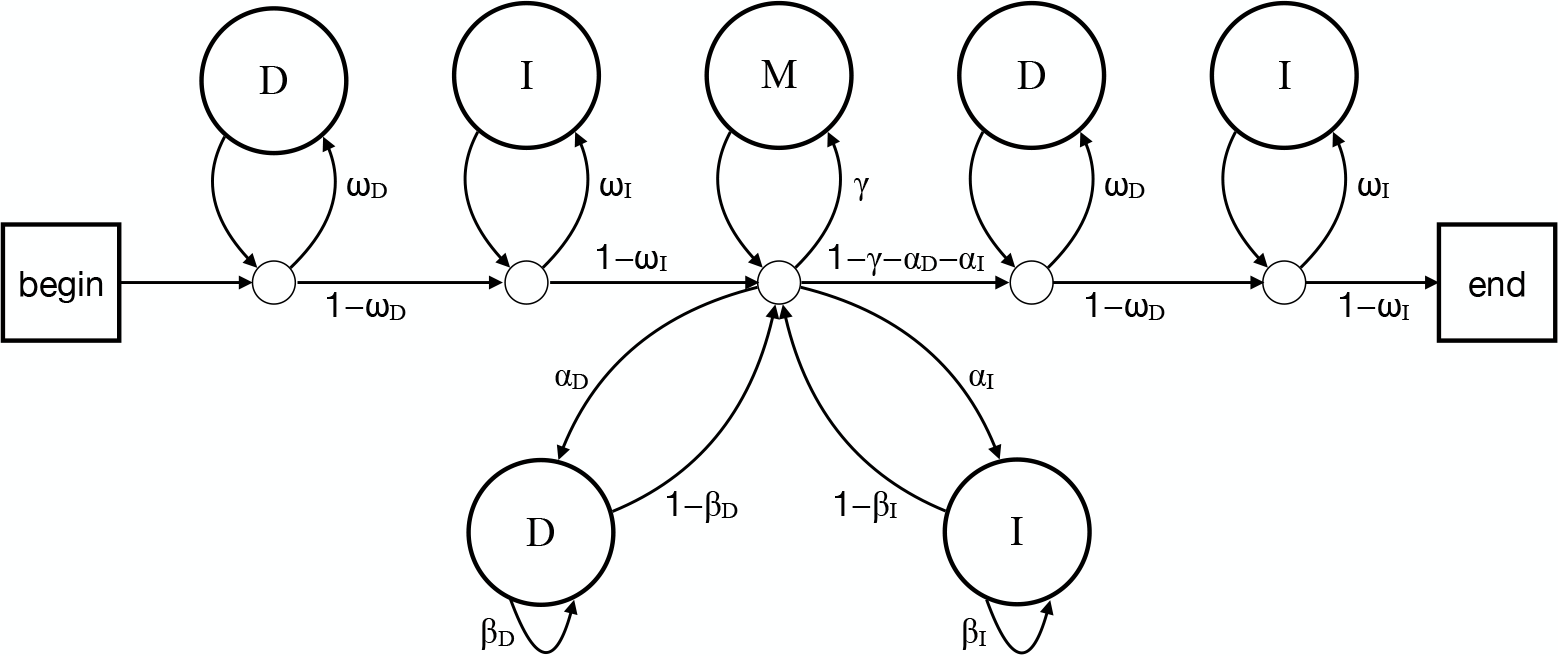
A probability model for two sequences with a related region. The model produces different alignments with different probabilities. Starting at “begin”, the arrows are followed according to their probabilities (e.g. *ωD* versus 1 − *ω*_*D*_ ). Each pass through a **D** state produces an unaligned letter *x* in one sequence (*R*), with probabilities *φ*_*x*_. Each pass through an **I** state produces an unaligned letter *y* in the other sequence (*Q*), with probabilities *ψ*_*y*_ . Each pass through **M** generates two aligned letters, *x* in *R* and *y* in *Q*, with probabilities *π*_*xy*_ .

Lunter et al. compared DNA sequences accurately, using probability methods similar to the present study [18]. In their scenario, they are given a pair of sequences that are assumed to have one related region. So they didn’t use any score to find related regions.

HMMER uses a log likelihood ratio similar to eq. 4 [4]. HMMER is more general, however. It finds related parts between a sequence and a sequence family: the family has position-specific monomer and gap probabilities. The null hypothesis (no related parts) is compared to either a “unihit” hypothesis (one related part), or a “multihit” hypothesis (one or more related parts). Each hypothesis has probabilities for the length of the sequence. The hypotheses are roughly equal in this regard. HMMER achieves that by adjusting each hypothesis to fit the length of each sequence that is provided for analysis [4].

FEAST is practical heuristic software for finding related sequence parts, and it sums probabilities of alternative alignments [19]. It doesn’t determine whether similarities would be unlikely to exist between random sequences. Compared to standard alignment, it is slower, and implementation difficulty was not made clear. It seems to be an implementation, rather than a simple method that can be widely implemented.

Several of these previous methods use a likelihood ratio, which is the optimal way to judge between two hypotheses with those likelihoods [20]. It is however not clear that comparing two hypotheses is the best way to find related sequence parts. For example, if we compare the human and mouse X chromosomes, it is not very useful just to know whether or not they have 0 related parts. There is a tradeoff between judging whether sequences are related, and how they are related. A single alignment specifies a relationship in detail, but has the lowest power to detect subtle relationships. Summing over all alignments has the highest power, but least detail. We seek a compromise between these extremes.

The final previous idea that we shall examine is “hybrid alignment” [21]. When hybrid alignment compares a sequence of length *m* to a sequence of length *n*, it considers *m* × *n* hypotheses for how they are related. Each hypothesis concerns the the length-*I* prefix of one sequence and length-*j* prefix of the other. The hypothesis is that the sequences have a related region that ends exactly at the ends of these prefixes. The probability associated with this hypothesis is the sum of probabilities of all alignments that start anywhere and end at the ends of the prefixes. This sum of alignment probabilities is converted to a score (eq. 2). Remarkably, it is easy to calculate the probability of an equal or higher score between random i.i.d. sequences, provided that the alignment parameters satisfy a “conservation condition”. This calculation was not proven correct, but was supported by theoretical arguments and tests on random sequences.

The present study builds on hybrid alignment. Hybrid alignment has been neglected, probably because its description was mathematically fearsome, and it was presented as a strange “hybrid” method. Here, instead of alignments ending at (*i, j*), we shall sum over alignments that pass through (*i, j*). This treats uncertainty about starts and ends of related regions symmetrically. It is a good fit for practical heuristic aligners, which extend alignment forwards and backwards from a candidate starting point.

## Methods

### Review of standard alignment

Practical methods for finding related parts of sequences often have two steps. To cope with big data, the first step uses fast heuristics to find potentially related regions, then the second step defines the regions precisely by alignment. The heuristics are various: short exact matches, spaced matches, reduced-alphabet matches, groups of nearby matches, etc. [7, 22–24]. The second step is often done by extending alignments to the left and right of a small “core” alignment (fig. 1B). This finds the highest-scoring alignment extension, within a limited range shown in gray in fig. 1B. The range is adjusted as the algorithm proceeds, attempting to follow the highest score. The best way to do this is unclear: several ways have been suggested [7, 25–28].

To avoid irrelevant complexity, here is the extension algorithm in a block of predefined size *n × n* (fig. 1A). This algorithm extends forwards (towards the lower-right in fig. 1A): a similar algorithm extends backwards. The two sequences are *R* (e.g. “reference”) and *Q* (e.g. “query”), with positions numbered as in fig. 1A. The score for aligning monomer *x* (a, c, g, or t for DNA) in *R* to monomer *y* in *Q* is *S*_*xy*_. Let us use different scores for starting versus extending gaps, because gaps are somewhat rare (low start probability) but often long (high extension probability). So the score for a length-*k* deletion is *a*_*D*_ + *b*_*D*_ × (*k* − 1), for a length-*k* insertion *a*_*I*_ + *b*_*I*_ × (*k* − 1).

#### Algorithm 1

Optimum alignment extension in an *n* × *n* block (lower-right gray block in fig. 1A)

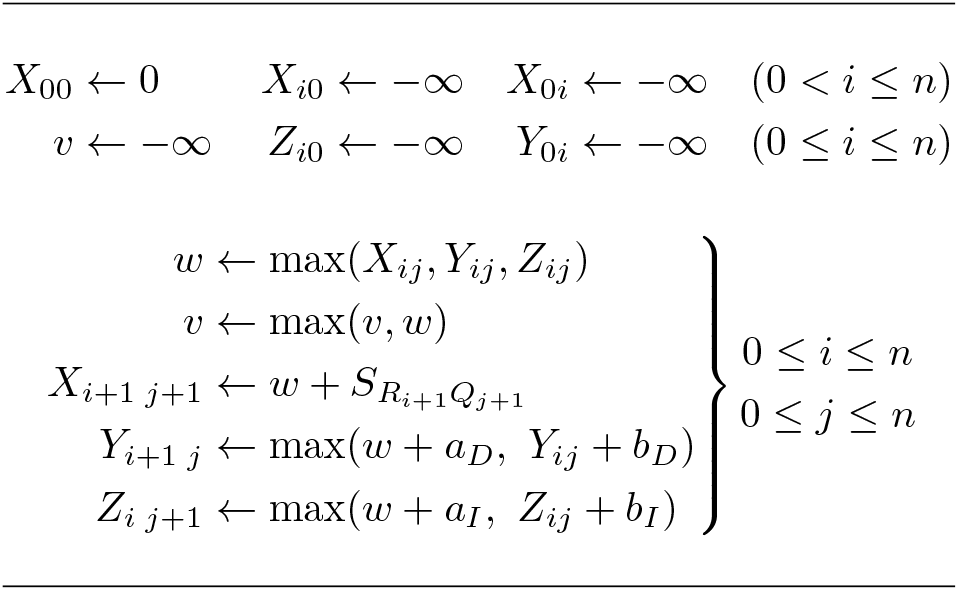

Alg. 1 finds the highest possible score for an alignment ending at *R*_*i*_ and *Q*_*j*_, with *R*_*i*_ aligned to *Q*_*j*_ (*X*_*ij*_), or *R*_*i*_ aligned to a gap (*Y*_*ij*_), or *Q*_*j*_ aligned to a gap (*Z*_*ij*_). *v* tracks the best alignment score seen so far, and is the final output. Several variants of this algorithm are possible. This variant has simple boundary conditions, while minimizing add, max, and array access operations [29]. The alignment score after left and right extension is:

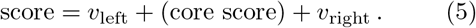

### New extension algorithm

This section shows the new algorithm, and the next section explains how it sums probabilities. There are two changes from alg. 1. The first, which makes no difference to the result, is to work with exponentiated values: probability ratios instead of log probability ratios. This is so we can easily add them. This change means that −∞ ⇒ 0, 0 ⇒ 1, and addition ⇒ multiplication. The other change is: maximization ⇒ addition. This calculates the sum of probability ratios for all alignment extensions in the gray area (eq. 2), instead of the maximum probability ratio. The score with left and right extension is:

#### Algorithm 2

Sum over alignment extensions in an *n × n* block (lower-right gray block in fig. 1A)

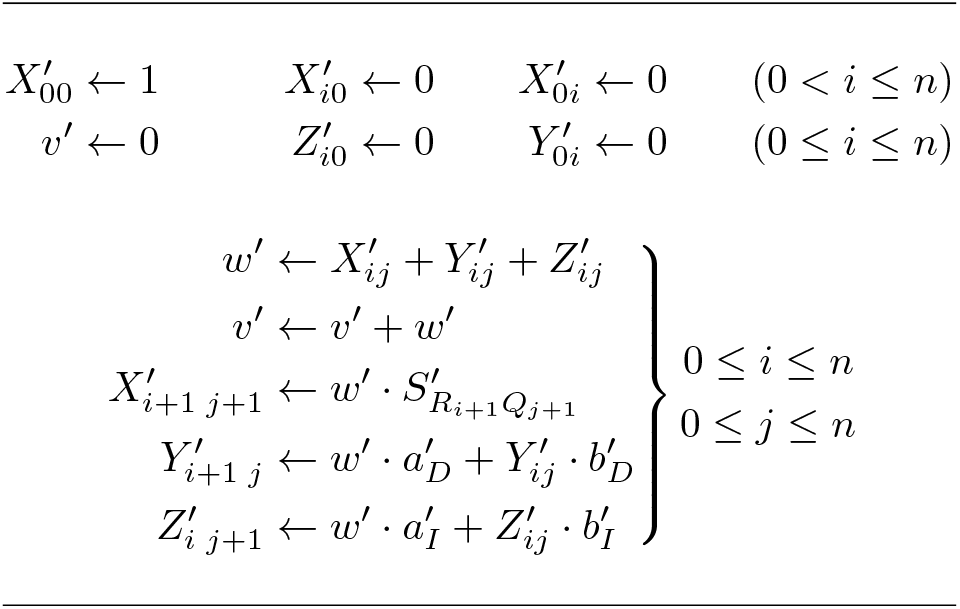

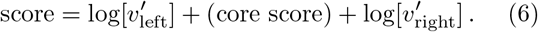

Note that the number of computational operations is unchanged: the only changes are replacing addition with multiplication, and maximization with addition. So the computational complexity is unchanged. The practical run time depends on implementer expertise (e.g. using SIMD), but plausibly it need not increase too much.

Unfortunately, the values in alg. 2 may become too large for the computer to handle. This can be fixed by occasionally rescaling them (see the Supplement). Because the rescaling is only done occasionally, it does not increase the run time noticeably.

### Parameters from probabilities

To use alg. 1 or 2, we must choose values for the parameters (e.g.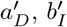). They are related to probabilities of matches, mismatches, and gaps. We need a precise definition of these probabilities. We shall use the definition in fig. 2. The main alignment probabilities are shown in the middle: *α*_*D*_ is the deletion start probability, *β*_*D*_ the deletion extension probability, *α*_*I*_ the insertion start probability, and *β*_*I*_ the insertion extension probability. Apart from starting an insertion or deletion, the other possibilities are to have a pair of aligned monomers (probability *γ*), or end the alignment (probability 1 − *γ* − *α*_*D*_ − *α*_*I*_ ). A pair of aligned monomers has probability *π*_*xy*_ of being monomer type *x* in sequence *R* and *y* in sequence *Q*. There are also probabilities for monomers left and right of the alignment: *ω*_*D*_ and *ω*_*I*_ (for sequence *R* and *Q* respectively). An unaligned monomer is of type *z* with probability *φ*_*z*_ in *R*, and *ψ*_*z*_ in *Q*.

A “null alignment” is produced by a path that never traverses the *α*_*D*_, *α*_*I*_, or *γ* arrows. This corresponds to a length-0 alignment between the sequences. The score of a null alignment is traditionally zero, which is achieved by eq. 1. This means that an alignment score is determined solely by the aligned regions, and not by flanking sequences or their lengths.

Having defined these probabilities, we can state the parameters for alg. 2, as shown previously [3]:

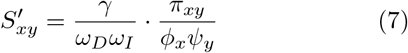

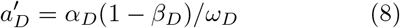

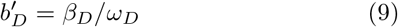

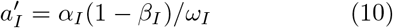

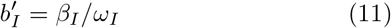

With these parameters, alg. 2 calculates the sum of probability ratios in eq. 2. The parameters for alg. 1 are defined similarly (see the Supplement).

Fig. 2 is not the only way to define the probabilities [3]. There are other ways that cause no change to alg. 1, but cause slight changes to alg. 2. The definition in fig. 2 seems to produce the simplest equations, and minimizes the number of computational operations in alg. 2. This is a minor improvement over hybrid alignment, which defined probabilities with an unnatural asymmetry between insertions and deletions [3, fig. S2].

### A non-heuristic similarity score

What would we do if we did not need fast heuristics? As mentioned in the Introduction, we seek a compromise between using just one alignment, and summing over all alignments. We shall use a compromise with two useful features: it is approximated by the heuristic approach (fig. 1B), and we can tell if a similarity is unlikely to exist between random sequences.

When we compare a length-*m* sequence (*R*_1_, *R*_2_, …, *R*_*m*_) to a length-*n* sequence (*Q*_1_, *Q*_2_, …, *Q*_*n*_), we shall consider (*m* + 1) *×* (*n* + 1) hypotheses for how they are related. Imagine cutting a length-*i* prefix of *R* and a length-*j* prefix of *Q*. These cuts define a point, illustrated in fig. 1C by the touching corners of the gray rectangles. The hypothesis is that the sequences are related by any alignment that passes through this point (including alignments that start or end there). The score of this hypothesis is given by eq. 2, with the sum taken over these alignments.

We can calculate this score for each (*i, j*). We first calculate 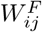, the sum of probability ratios of all alignments that end immediately after *i* letters of *R* and *j* letters of *Q*:

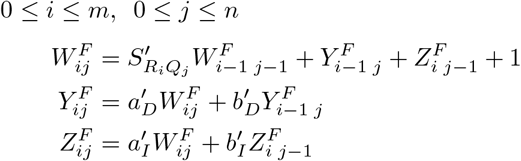

The boundary condition is: when *i <* 0 or *j <* 0, 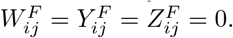.

We then calculate 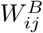, the sum of probability ratios of all alignments that start immediately after *i* letters of *R* and *j* letters of *Q*:

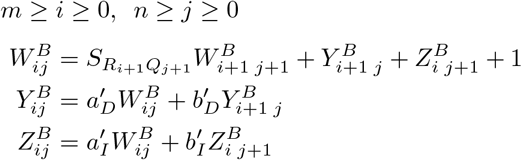

The boundary condition is: when *i > m* or *j > n*,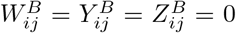. Finally, the sum of probability ratios of all alignments passing through (*i, j*) is 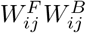. So the score is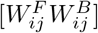.

A minor point is that this calculation excludes alignments that pass through (*i, j*) while traversing the *β*_*D*_ or *β*_*I*_ arrow (fig. 2). It’s not clear whether it’s better to include them, but it seems unlikely to make much difference.

### Similarity scores occurring by chance

We wish to know the probability of getting a similarity score, 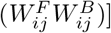, between random sequences. Because our score is similar to that of hybrid alignment [21], we shall assume (and test) that we can use the same approach. This approach requires that the alignment parameters satisfy a “conservation condition”.

The conservation condition is related to alignment length bias. The probabilities in fig. 2 may be biased towards short or long alignments between two given sequences. For example, if *α*_*D*_, *α*_*I*_, and *γ* are all low (almost 0), but *ω*_*D*_ and *ω*_*I*_ are high (almost 1), shorter alignments will have higher probability. In the converse situation, longer alignments are favored. It was shown previously [3] that the probabilities are unbiased when:

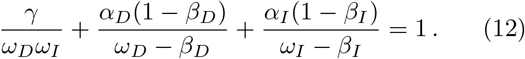

Eq. 12 is a special case of the conservation condition. The general case is shown in the Appendix, but we shall only use the special case.

We can calculate the probability of a similarity score between two random i.i.d. sequences: one with length *m* and monomer probabilities *φ*_*x*_, the other with length *n* and probabilities *ψ*_*y*_. These monomer probabilities must match those in eq. 7. The prediction is that

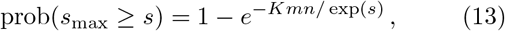

in the limit where *m* and *n* are large. exp is defined to mean inverse of log.

Eq. 13 has one unknown parameter, *K*. We can estimate *K* by generating some pairs of random i.i.d. sequences, calculating *s*_max_ for each pair, and fitting *K*. The maximum likelihood fit is:

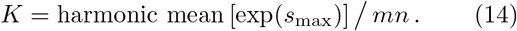

The random chance of a similarity score *s* is often reported as an *E*-value. The *E*-value is the expected number of similarities with score ≥ *s* between two random sequences of length *m* and *n*. Chance similarities occur at a constant rate between any part of one sequence and any part of the other. So they follow a Poisson distribution. This means the *E*-value is:

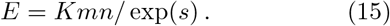

### Implementation and availability

This approach to finding related sequence parts was added to LAST (https://gitlab.com/mcfrith/last). This implementation is just a proof of principle: the heuristics are not necessarily the best. LAST can use either alg. 1 with *E*-values from the ALP library, or alg. 2. In both cases, it gets a representative alignment by running alg. 1 and tracing back a way to get the maximum *v*. LAST runs these algorithms not in an *n × n* block (fig. 1A), but in a range that is adjusted as the algorithm proceeds (fig. 1B, see the Supplement).

LAST estimates *K* from random sequences. It is not clear how long these sequences need to be. Based on some tests (see the Supplement), LAST uses 50 pairs of length-500 sequences by default.

The match/mismatch/gap probabilities (*π*_*xy*_, *α*_*D*_, etc.) were determined using `last-train`, which finds related regions of given sequences and infers these probabilities [30]. `last-train` was modified to make the probabilities satisfy eq. 12. It sets *ω*_*D*_ and *ω*_*I*_ by assuming that they are equal, and finding the unique value that satisfies eq. 12 (see the Appendix). `last-train` is not necessarily the best way to get the probabilities. Another option is to use BLOSUM or PAM substitution probabilities (*π*_*xy*_), which are in the NCBI C`++` toolkit (version 25.2, in `matrix frequency data.c`).

The `last-train` output files from this study are at https://gitlab.com/mcfrith/sum-align. There too is a simple Python script to help use these ideas. It takes as input *α*_*D*_, *α*_*I*_, *β*_*D*_, *β*_*I*_, and *γ*, finds *ω*_*D*_ = *ω*_*I*_ that satisfies eq. 12, and outputs 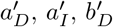, and 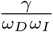 . LAST includes code for calculating the non-heuristic similarity score (used for estimating *K*), in a self-contained file (`mcf alignment path adder.cc`).

## Results

### Similarity scores occurring by chance

Let us test whether we can calculate probabilities of similarity scores between random sequences. Three sequence-search scenarios are considered, with typical DNA, `at`-rich DNA, and proteins.

The first scenario is finding ancient (shared with reptiles) repeats in the human genome. This can be done by comparing the genome to repeat consensus sequences from Dfam [31]. Some are relics of transposable elements, others are source unknown.

The match/mismatch/gap probabilities were found by comparing the genome to the consensus sequences with `last-train`. Then, pairs of random i.i.d. sequences were generated, with monomer probabilities *φ*_*x*_ and *ψ*_*y*_ equal to those found by `last-train`. For each pair of sequences, the highest score *s*_max_ was found by the non-heuristic algorithm. The observed scores are consistent with their probabilities predicted by eq. 13 (fig. 3 left).

**Figure 3:**
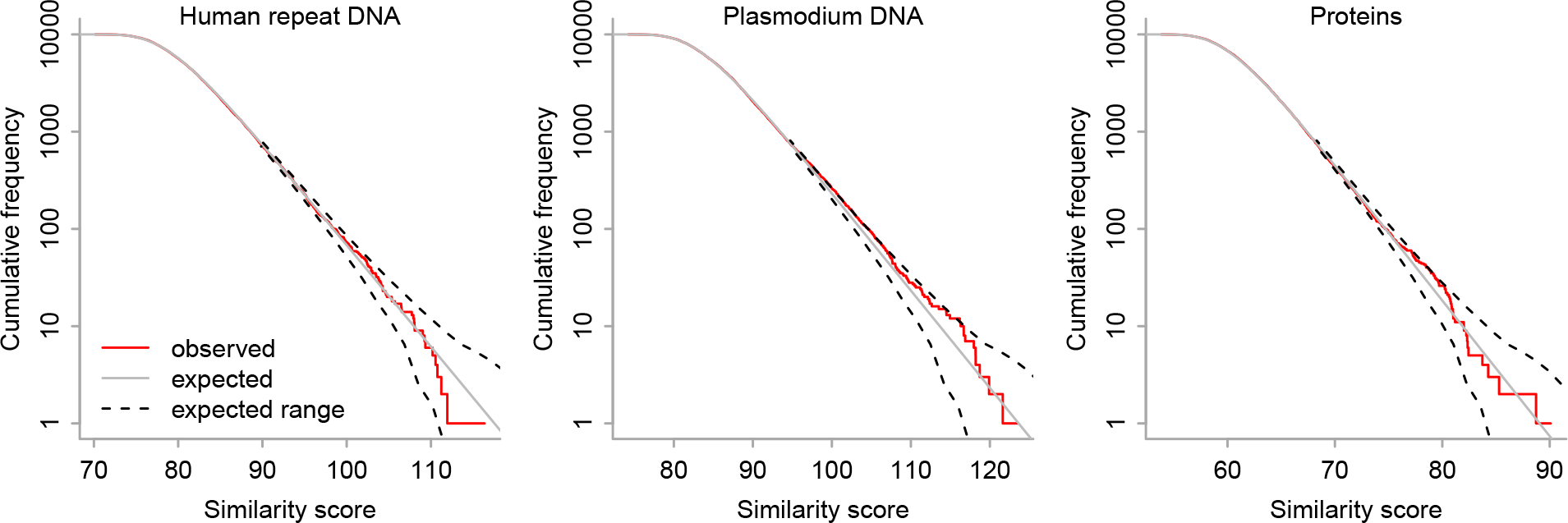
Non-heuristic similarity score (*s*_max_) between 10000 pairs of random i.i.d. sequences (length 10000), for three sets of alignment parameters. The frequency has a 5% chance of being outside the dashed lines (2.5% each).

The next scenario was to find related parts of the *Plasmodium falciparum* and *Plasmodium yoelii* genomes, which are both about 80% `a`+`t`. Alignment parameters were found by `last-train`, then similarity scores were found between random `at`-rich sequences (fig. 3 middle). Again, the observed scores are consistent with eq. 13.

The third scenario was to find related parts of proteins from *Aquifex aeolicus* (a heat-loving bacterium) and *Pyrolobus fumarii* (a heat-loving archaeon). Alignment parameters were found by `last-train`, then similarity scores were found between random sequences (fig. 3 right). Yet again, the observed scores are consistent with eq. 13.

### Related versus biased sequences

A high similarity score could occur because the sequence regions have a common ancestor, and/or similar composition bias (fig. 4A). We can investigate this by comparing two biological sequences after reversing (but not complementing) one of them (but see [32]). Then there are no segments related by evolution, but there may be similarities due to composition bias (fig. 4A).

**Figure 4:**
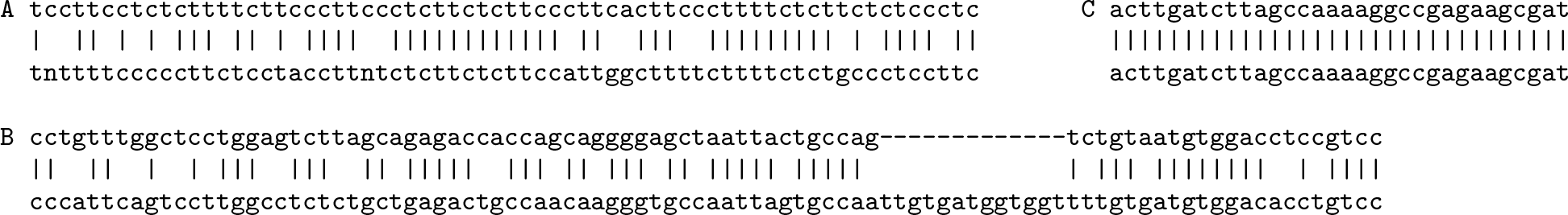
Alignment between parts of: **A** reversed human chromosome Y (upper), and UCON1 (lower). This is the highest-scoring alignment in the top-left panel of fig. 5. **B** Human chromosome 6 (upper) and LFSINE Vert (lower). **C** Human chromosome 21 (upper) and human U2 (lower).

Thus, the reversed human genome was searched against the repeat consensus sequences, the reversed *P. yoelii* genome against *P. falciparum*, and reversed *P. fumarii* proteins against *A. aeolicus* proteins. In each case, alg. 2 found more high similarity scores than expected between random sequences (fig. 5 upper row), due to biased regions (fig. 4A).

**Figure 5:**
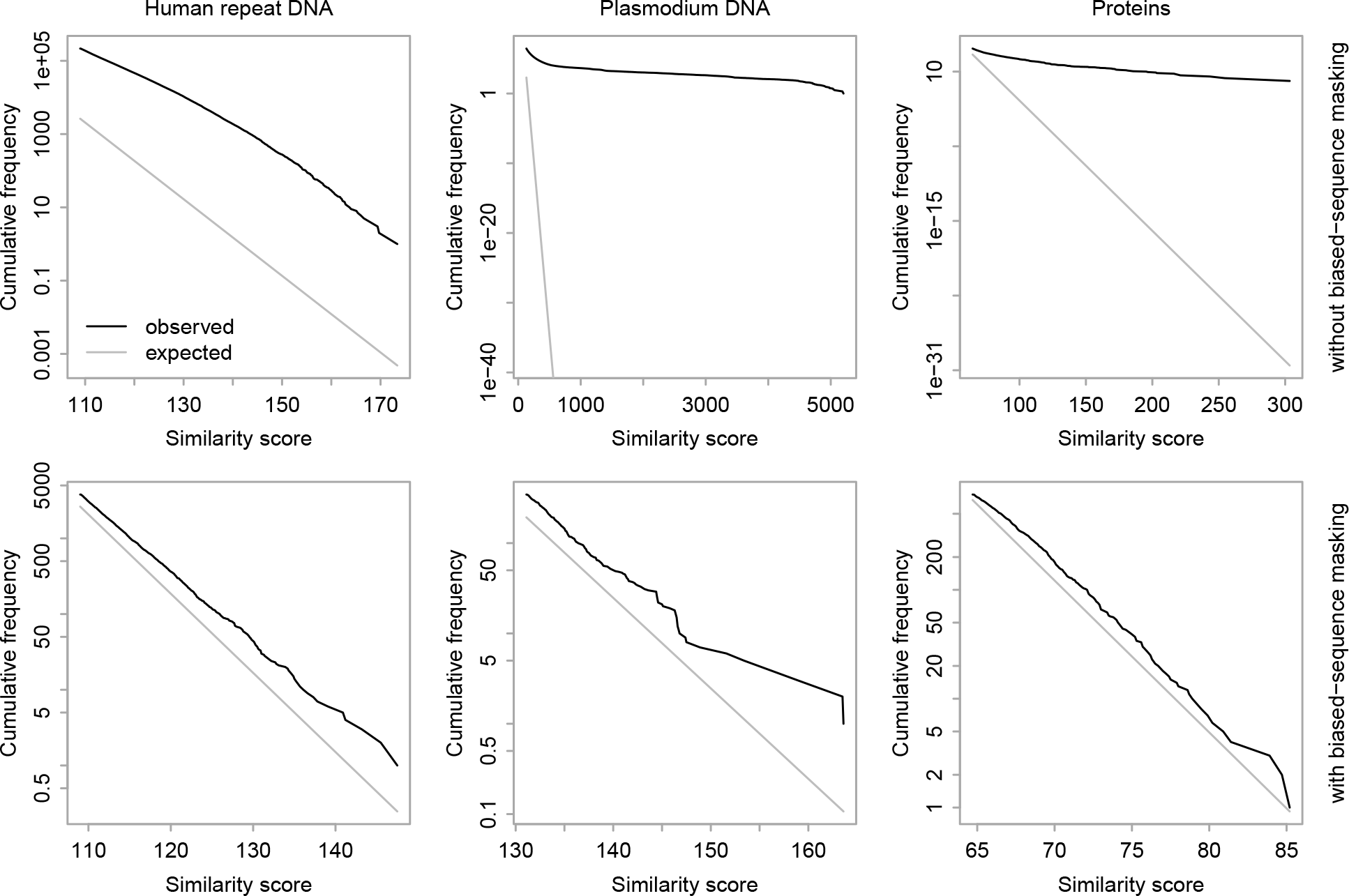
Heuristic similarity scores, between reversed and non-reversed sequences, with or without biased-sequence masking.

For standard alignment, these similarities can be avoided by detecting biased regions with `tantan` [33], and “masking” them. An effective masking scheme is to change *S*_*xy*_ ← min(*S*_*xy*_, 0) when *x* or *y* is in a biased region [34]. For alg. 2, this masking scheme was tried: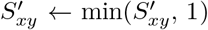 .^1^ With this masking scheme, the similarity scores between reversed and non-reversed sequences are close to what is expected between random sequences (fig. 5 lower row).

### The new method can be more sensitive

Related sequence parts were sought by standard alignment (alg. 1) and by the new method (alg. 2), with biased-sequence masking. The two methods often find identical alignments, but with different scores and *E*-values. The new method tended to assign lower *E*-values, for alignments of the human genome to repeat consensus sequences (fig. 6A), and *P. fumarii* proteins to *A. aeolicus* proteins (fig. 6B). For example, both methods found the same alignment between human chromosome 6 and the LFSINE Vert consensus (fig. 4B): the alg. 1 *E*-value is 22, and the alg. 2 *E*-value is 0.000031. This means the new method can find related sequence parts more confidently.

**Figure 6:**
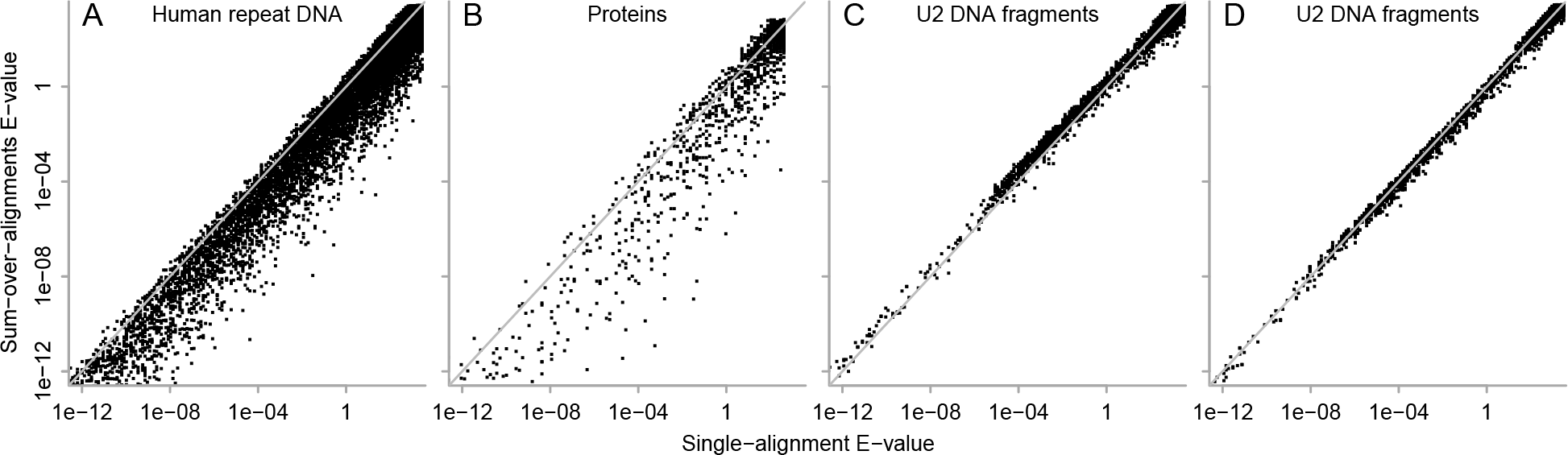
*E*-values for identical alignments found by alg. 1 (horizontal axis), and alg. 2 (vertical axis). Each point is one alignment. The diagonal gray lines indicate equal *E*-values. These *E*-values were calculated with *m* = total length of all reference sequences (e.g. all *A. aeolicus* proteins), and *n* = total length of all query sequences (e.g. all *P. fumarii* proteins). For DNA, one query strand was searched against both reference strands, so *m* was multiplied by 2.

In other words, at a fixed *E*-value cutoff, the new method predicts more related regions. The preceding results indicate that the *E*-values are not overconfident. This implies that the new method finds more true positives for a given false positive rate.

### The new method can be less sensitive

Another sequence search scenario was considered: finding U2 fragments in the human genome. U2 is one of the small nuclear RNAs that splice introns. The genome has many fragmentary copies of it, which are thus considered repeats.

The human genome was searched against a U2 DNA sequence, using the match/mismatch/gap probabilities found earlier for the human genome and ancient repeat consensus sequences. This time, alg. 2 tended to assign higher *E*-values than alg. 1 (fig. 6C). For example, both methods found the alignment shown in fig. 4C: the alg. 1 *E*-value is 0.00021, and the alg. 2 *E*-value is 0.0011.

The likely explanation is as follows. The genome has many short, near-exact copies of the first 30–40 bases of U2. The match/mismatch/gap probabilities, however, reflect greater sequence divergence (e.g. fig. 4B). These mistuned probabilities pessimize both algorithms, but they pessimize alg. 2 more. That is because probabilities with greater divergence attach more weight to alternative alignments.

This suggests that alg. 2 should work better than alg. 1 if the probabilities are a good fit to the sequences. To test this, match/mismatch/gap probabilities were found by comparing the genome to the U2 sequence with `last-train`. Then, the genome was searched against U2 using these probabilities. This time, alg. 2 tends to assign slightly lower *E*-values than alg. 1 (fig. 6D). Even with well-tuned probabilities, the advantage of alg. 2 is expected to decrease for more similar sequences, because the evidence for relatedness is more concentrated in one alignment.

## Discussion

We have seen how to find related parts of sequences, by summing probabilities of alternative alignments between them. This is better than standard alignment at detecting subtle relationships, at least in a few tests. It can be worse than standard alignment, when the probabilities are tuned for high divergence but the sequences have low divergence. The new method is more beneficial for higher divergence, where alternative alignments contribute more. In a spectacular example, it found gene-regulating DNA conserved in diverse animals since the Precambrian [35].

A major further benefit is that the new method simplifies *E*-value calculation. This makes it easier to experiment with different alignment parameters, beyond the BLAST parameter sets. This can be useful for proteins [36], but especially for DNA, because there has been less effort to optimize DNA parameters.

Moreover, the new method can be generalized to other kinds of alignment. For example, we used the same approach for DNA-versus-protein alignment [37]. We defined probabilities similarly to fig. 2, but with extra probabilities for frameshifts. We summed probabilities of alternative alignments, and accurately predicted similarity scores between random sequences. This method was highly sensitive: it found ancient and degraded relics of Paleozoic mobile elements in vertebrate genomes. Some previous methods have used two kinds of alignment gap: short-and-frequent and long-and-rare [15, 18]. This can be done by elaborating fig. 2. The approach of this paper is predicted to work in that case too. This makes it easier to try such different kinds of alignment, in particular because it provides *E*-values. Another alignment elaboration is position-specific match, mismatch, and gap rates [2, 4, 38, 39]: this was incorporated into hybrid alignment [40].

By combining alg. 2 with another similar algorithm (see the Supplement), we can calculate the probability that each alignment column is correct (fig. 7), based on the assumed rates of matches, mismatches, and gaps [2, 15]. It is also possible to make alignments based on these probabilities: for example, align monomers whose alignment probability is *>* 0.5 [41].

**Figure 7:**
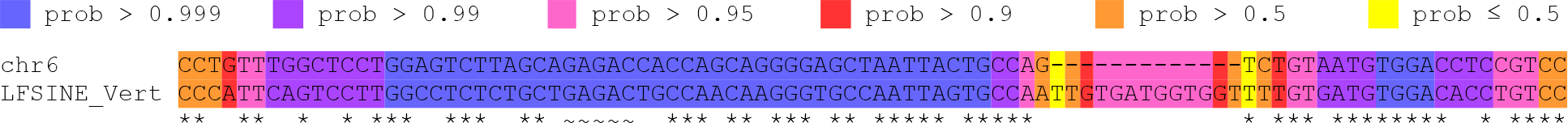
An alignment (the same as fig. 4B) with colors indicating the probability that each column is correct. _ indicates the “core” (fig. 1), whose column probabilities are assumed to be 1. *\** indicates non-core exact matches.

For short sequences, the *E*-values are too high. Eq. 15 assumes the number of chance similarities is proportional to *mn*. That becomes too high for short sequences, because a similarity must have non-zero length and fit in the sequences. There are corrections for this effect, including in hybrid alignment, but it is somewhat complex [21].

We should consider some basic aims and assumptions. Suppose we seek related regions between a genome of length *m*, and *q* “query” sequences with lengths *n*_1_, *n*_2_, …, *n*_*q*_. We could assume that related regions are equally likely to occur per query sequence, or per unit sequence length. We could aim to find relationships for as many query sequences as possible, or as many related regions as possible. If part of query *i* is similar to part of the genome, we can calculate an *E*-value from eq. 15 with sequence lengths *m* and *n*_*i*_. This increasingly disfavors related regions in increasingly long queries. As mentioned in the Introduction, some previous methods similarly penalize related regions in longer sequences. The alternative is to use eq. 15 with sequence lengths *m* and 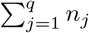.

This treats related regions equally, regardless of the containing sequence’s length. The aim is important when testing sequence search methods, because the test may count the number of sequences or related regions found.

## Acknowledgments

I am grateful to John Spouge for encouragement that the *E*-values would be simple, Yi-Kuo Yu for advice about hybrid alignment, and Travis Wheeler for advice about HMMER.

## Appendix

### Conservation condition

Suppose we compare two random i.i.d. sequences, with monomer probabilities Φ_*x*_ and Ψ_*y*_. The conservation condition is In the special case where Φ_*x*_ = *φ*_*x*_ and Ψ_*y*_ = *ψ*_*y*_, eq. 16 is the same as eq. 12.

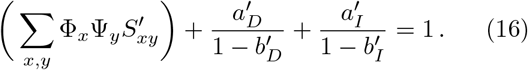

The conservation condition depends on the precise definition of gap probabilities (fig. 2), as explained previously [3]. The conservation condition for hybrid alignment was different [21, eq. 28], because they defined gap probabilities differently [3, fig. S2].

The conservation condition generalizes an equation in the proven *E*-value solution for gapless alignment [6, eq. 4].

### Finding *ω*_*D*_ = *ω*_*I*_ that satisfies eq. 12

First, assume that *ω*_*D*_ *> β*_*D*_ and *ω*_*I*_ *> β*_*I*_ . Otherwise, alignment gaps would have non-negative scores, which is not useful for sequence comparison.

As *ω*_*D*_ = *ω*_*I*_ increases from max(*β*_*D*_, *β*_*I*_ ) to 1, the LHS of eqn. 12 decreases, from a value *>* 1 to a value *<* 1. So it equals 1 at a unique point, which can be found by e.g. repeated bisection.

#### prob(null alignment)

If we define gap probabilities as in fig. 2, a null alignment is produced by a path that never traverses the *α*_*D*_, *α*_*I*_, or *γ* arrows. Suppose we compare a length-*m* sequence *R* to a length-*n* sequence *Q*.

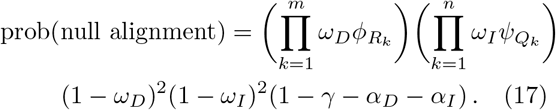

#### Footnotes

1 Another untested idea is 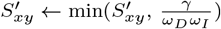.

