## Supplement for "A simple theory for finding related sequences by adding probabilities of alternative alignments"

Martin C. Frith

April 13, 2024

#### Rescaling to avoid overflow in Algorithm 2

Here is one way to avoid overflow, used in this study's implementation. First, initialize an extra variable: $u \leftarrow 0$. Then, execute alg. 2 in order of increasing antidiagonal: $i + j$. Just after calculating every 16th antidiagonal, update $u$:

$$u \leftarrow u + \log(v') \,. \tag{S1}$$

Rescale all the $X$, $Y$, and $Z$ values that were calculated on this and the previous antidiagonal:

$$X'_{ij} \leftarrow X'_{ij}/v' \tag{S2}$$

$$Y'_{ij} \leftarrow Y'_{ij}/v' \tag{S3}$$

$$Z'_{ij} \leftarrow Z'_{ij}/v' \tag{S4}$$

Then, reset $v' \leftarrow 1$. After the algorithm finishes, the final extension score is $u + \log(v')$.

#### $x$-drop implementation

LAST defines regions in which it seeks alignment extensions (gray area in fig. 1B) by an $x$-drop method, similar to one described previously [1]. For alg. 1, it executes the algorithm in order of increasing antidiagonal: $k = i + j$. When executing alg. 1 on antidiagonal $k$ (i.e. when $i + j = k$), $i$ is restricted to a range: $A_k \le i \le B_k$. It starts with $A_0 = B_0 = 0$, and adjusts $A_k$ and $B_k$ as follows:

$$A_{k+1} = \begin{cases} A_k & \text{if } W_{A_k\, k-A_k} \ge V_k - x \\ A_k + 1 & \text{otherwise} \end{cases} \tag{S5}$$

$$B_{k+1} = \begin{cases} B_k + 1 & \text{if } W_{B_k\, k-B_k} \ge V_k - x \\ B_k & \text{otherwise} \end{cases} \tag{S6}$$

Here, $W_{ij}$ is defined to be $\max(X_{ij}, Y_{ij}, Z_{ij})$. Also, $V_k$ is the value of $v$ (the highest score seen so far) just before executing alg. 1 on antidiagonal $k$. $x$ is the maximum score drop. The recommended value for $x$ is just below the minimum score for reporting an alignment [2, 3].

For alg. 2, LAST first runs alg. 1 and then re-uses the same gray area for alg. 2. The best method is unclear. LAST's DNA-versus-protein alignment is different: it's version of alg. 2 has its own $x$-drop-like algorithm [4]. If we do not re-use the same gray area, the similarity score (from alg. 2) may be inconsistent with the representative alignment (from alg. 1), e.g. huge score but tiny alignment.

### Parameters for Algorithm 1

The parameters for alg. 1 are defined like this [5, Supplement 3.1]:

$$S_{xy} = \log[S'_{xy}] \tag{S7}$$

$$a_D = \log[a'_D] \tag{S8}$$

$$a_I = \log[a'_I] \tag{S9}$$

$$b_D = \log[b'_D + a'_D] \tag{S10}$$

$$b_I = \log[b'_I + a'_I] \tag{S11}$$

Eq. S10 defines the probability of extending a deletion as the sum of two probabilities: extension via the $\beta_D$ arrow, and extension via the $1 - \beta_D$ and $\alpha_D$ arrows (fig. 2). Eq. S11 does the same for insertions.

### Alignment column probabilities

We can calculate the probability that each pair of monomers is aligned. In other words, the probability that $R_i$ is aligned to $Q_j$. First, note that alg. 2 calculates $X'_{ij}$, which is the probability ratio sum of alignment extensions ending with $R_i$ aligned to $Q_j$. Another "backward" algorithm (alg. S1) calculates $W''_{ij}$: the probability ratio sum of alignments starting just after $R_i$ aligned to $Q_j$. This means that $X'_{ij}W''_{ij}$ is the probability ratio sum of all alignment extensions that include $R_i$ aligned to $Q_j$. Finally,

$$\text{prob}(R_i \text{ aligns to } Q_j) = X'_{ij}W''_{ij}/v' \,. \tag{S12}$$

Here, $v'$ is the output of alg. 2: the probability ratio sum of all alignment extensions.

It's also possible to calculate gap probabilities:

$$\text{prob}(R_i \text{ aligns to a gap between } Q_j \text{ and } Q_{j+1}) = Y'_{ij}Y''_{ij}/v' \,, \tag{S13}$$

$$\text{prob}(Q_i \text{ aligns to a gap between } R_j \text{ and } R_{j+1}) = Z'_{ij}Z''_{ij}/v' \,. \tag{S14}$$

**Algorithm S1** Probability ratios of alignments starting at each point in an $n \times n$ block

$$W''_{n+1\,n+1} \leftarrow 0 \qquad W''_{i\,n+1} \leftarrow 0 \qquad W''_{n+1\,i} \leftarrow 0 \qquad (0 < i \leq n)$$
$$Z''_{i\,n+1} \leftarrow 0 \qquad Y''_{n+1\,i} \leftarrow 0 \qquad (0 \leq i \leq n)$$

$$\left.\begin{aligned} W''_{ij} &\leftarrow W''_{i+1\,j+1} \cdot S'_{R_{i+1}Q_{j+1}} + Y''_{i+1\,j} \cdot a'_D + Z''_{i\,j+1} \cdot a'_I + 1 \\ Y''_{ij} &\leftarrow W''_{ij} + Y''_{i+1\,j} \cdot b'_D \\ Z''_{ij} &\leftarrow W''_{ij} + Z''_{i\,j+1} \cdot b'_I \end{aligned}\right\} \begin{aligned} n \geq i \geq 0 \\ n \geq j \geq 0 \end{aligned}$$

The colors of gap columns in fig. 7 indicate the probability of aligning to a gap anywhere. In other words, $\sum_j Y'_{ij} Y''_{ij} / v'$ and $\sum_i Z'_{ij} Z''_{ij} / v'$.

It may be more efficient to replace the red 1 in alg. S1 with $1/v'$. This causes all the values calculated by the algorithm to be divided by $v'$, saving the division in eqs. S12–S14.

There are various ways to make alignments based on these probabilities. Two such ways are implemented in LAST: $\gamma$-centroid alignment and LAMA alignment [6]. These were not used in the present study.

### How the results were obtained

#### Sequence data

The genomes, proteins, and U2 DNA sequence (`NR_002716.3`) are from NCBI:

| Organism | Sequence data identifier |
|---|---|
| *Homo sapiens* | `hg38_no_alt_analysis_set` |
| *Plasmodium falciparum* | `GCF_000002765.5` |
| *Plasmodium yoelii* | `GCF_900002385.2` |
| *Aquifex aeolicus* | `GCF_000008625.1` |
| *Pyrolobus fumarii* | `GCF_000223395.1` |

The consensus sequences of ancient repeats (shared by mammals and reptiles) are at https://gitlab.com/mcfrith/sum-align. They are repeats from Dfam 3.7 with taxon "Amniota" and classification
`root;Interspersed_Repeat;Transposable_Element`
or
`root;Interspersed_Repeat;Unknown`.
Some repeats were excluded, because they are confusingly similar to younger repeats: `LINE/L2`, `LINE/CR1`, `MIR1_Amn`, `AmnSINE1`, `Chompy-6_Croc`, and `LmeSINE1c`.

#### Finding ancient repeats in the human genome

LAST version 1471 was used throughout. First, an index (called `repDB`) was made of the repeat consensus sequences:

```
lastdb -c -S2 -uMAM8 repDB repeats.fasta
```

- `-c` masks sequence regions that have biased composition.
- `-S2` indexes both DNA strands (so we needn't use both strands of the genome).
- `-uMAM8` increases sensitivity but also run time and memory use [7].

Next, rates of matches, mismatches, and gaps were found between the repeat sequences and the human genome:

```
last-train --revsym -X1 --sample-number=5000 -D1e9 repDB human.fasta > rep.train
```

- `--revsym` makes the match and mismatch rates strand-symmetric, e.g. the `a`:`c` rate equals the `t`:`g` rate.
- `-X1` treats unknown "n" bases in the repeats, which are numerous, as ambiguous: $S_{xy} = \log[\gamma/(\omega_D \omega_I)]$ if $x =$ `n`.
- `--sample-number=5000` increases the number of random genome fragments that `last-train` uses, for fear that the ancient repeats are rare in the genome.

- For the same reason, `-D1e9` sets a stricter similarity threshold. It uses similarities that are expected by chance $\leq$ once per $10^9$ genome base-pairs. (This is no longer recommended for LAST versions > 1519, because `last-train` tunes its `D` automatically.)

The reversed genome was searched against the repeats like this:

```
lastal -u3 -m100 -p rep.train -J1 repDB human-rev.fasta > rev.maf
```

- `-u3` applies masking throughout. (By default, after related regions have been found with masking, they are aligned without masking.)
- `-m100` makes it more sensitive and slow.
- `-J1` specifies alg. 2.

The (non-reversed) genome was searched against the repeats like this:

```
lastal -u3 -m100 -p rep.train -J$J -K1 repDB human.fasta > fwd.maf
```

- `$J` was replaced with either `0` (alg. 1) or `1` (alg. 2).
- `-K1` omits any alignment whose genome range lies in a higher-scoring alignment.

#### *Plasmodium* DNA analysis

First, an index (called `falcDB`) was made of the *P. falciparum* genome:

```
lastdb -c -R02 falcDB falciparum.fasta
```

- `-R02` uses `tantan` parameters suitable for `at`-rich DNA [8].

Then, rates of matches, mismatches, and gaps were found:

```
last-train --revsym falcDB yoelii.fasta > plasmo.train
```

The reversed *P. yoelii* genome was searched like this:

```
lastal -u3 -m100 -D1e5 -p plasmo.train -J1 falcDB yoelii-rev.fasta > rev.maf
```

#### Proteins

First, an index (called `aquiDB`) was made of the *A. aeolicus* proteins:

```
lastdb -c -p aquiDB Aquifex-prot.fasta
```

- `-p` specifies that these are protein sequences.

Then, rates of matches, mismatches, and gaps were found:

```
last-train -m1000 aquiDB Pyrolobus-prot.fasta > prot.train
```

The reversed *P. fumarii* proteins were searched like this:

```
lastal -u3 -m1000 -D1e3 -p prot.train -J1 aquiDB Pyrolobus-prot-rev.fasta > rev.maf
```

The (non-reversed) *P. fumarii* proteins were searched like this:

```
lastal -u3 -m1000 -D1e3 -p prot.train -J$J -K1 aquiDB Pyrolobus-prot.fasta > fwd.maf
```

## U2

First, an index (called `U2db`) was made of the U2 DNA sequence:

```
lastdb -c -S2 U2db U2.fasta
```

Then, rates of matches, mismatches, and gaps were found:

```
last-train --revsym --sample-number=50000 -D1e9 U2db human.fasta > U2.train
```

The human genome was searched against U2 like this:

```
lastal -u3 -p rep.train -J$J -K1 U2db human.fasta > fwd.maf
```

The two U2 results were obtained by running lastal with either `rep.train` or `U2.train`.

#### Similarity search without masking biased composition

Unmasked similarities were found by running `lastdb` without `-c` and `lastal` without `-u3`.

#### Length-dependence of random similarity scores

The formula for probability of similarity scores between random sequences (eq. 13) is inaccurate for short sequences. To check this, the non-heuristic similarity score ($s_{\max}$) was calculated for pairs of random i.i.d. sequences, of varying length (fig. S1). These scores were used to estimate $K$ (eq. 14). For sequence length 40 (fig. S1 top row), the similarity scores clearly differ from the expected distribution. The estimate of $K$ seems to stabilize by sequence length 10000, though it is impossible to prove that by such tests. For sequence length 500, the estimates of $K$ are too low by 8–16%.

#### Random match/mismatch similarity scores

It may be interesting to see similarity scores between random sequences, for the simplest possible alignment probabilities: gapless match/mismatch probabilities. This means that $\alpha_D = \alpha_I = 0$, and $\pi_{xy}$ has a fixed value when $x = y$, and a second fixed value when $x \neq y$. Four probability settings were checked (which correspond to simple match/mismatch scores):

| $\pi_{xy}$ $(x = y)$ | $\pi_{xy}$ $(x \neq y)$ | % identity | match score | mismatch score |
|---|---|---|---|---|
| 0.222 | 0.009337 | 89 | 2 | -3 |
| 0.1875 | 0.02083 | 75 | 1 | -1 |
| 0.1628 | 0.02905 | 65 | 5 | -4 |
| 0.141 | 0.03634 | 56 | 3 | -2 |

The non-heuristic similarity score ($s_{\max}$) was calculated for pairs of random i.i.d. sequences (fig. S2). These scores were used to estimate $K$ (eq. 14). Interestingly, $K$ increases as the match probability ($\pi_{xx}$) decreases. This is opposite to the behavior of $K$ for ordinary gapless alignment [9], but consistent with previous results for hybrid alignment [4, 10].

Another trend is that the estimate of $K$ converges more slowly as the match probability decreases. In other words, the % difference between $K$ estimated with sequence length 500 versus 10000 becomes bigger.

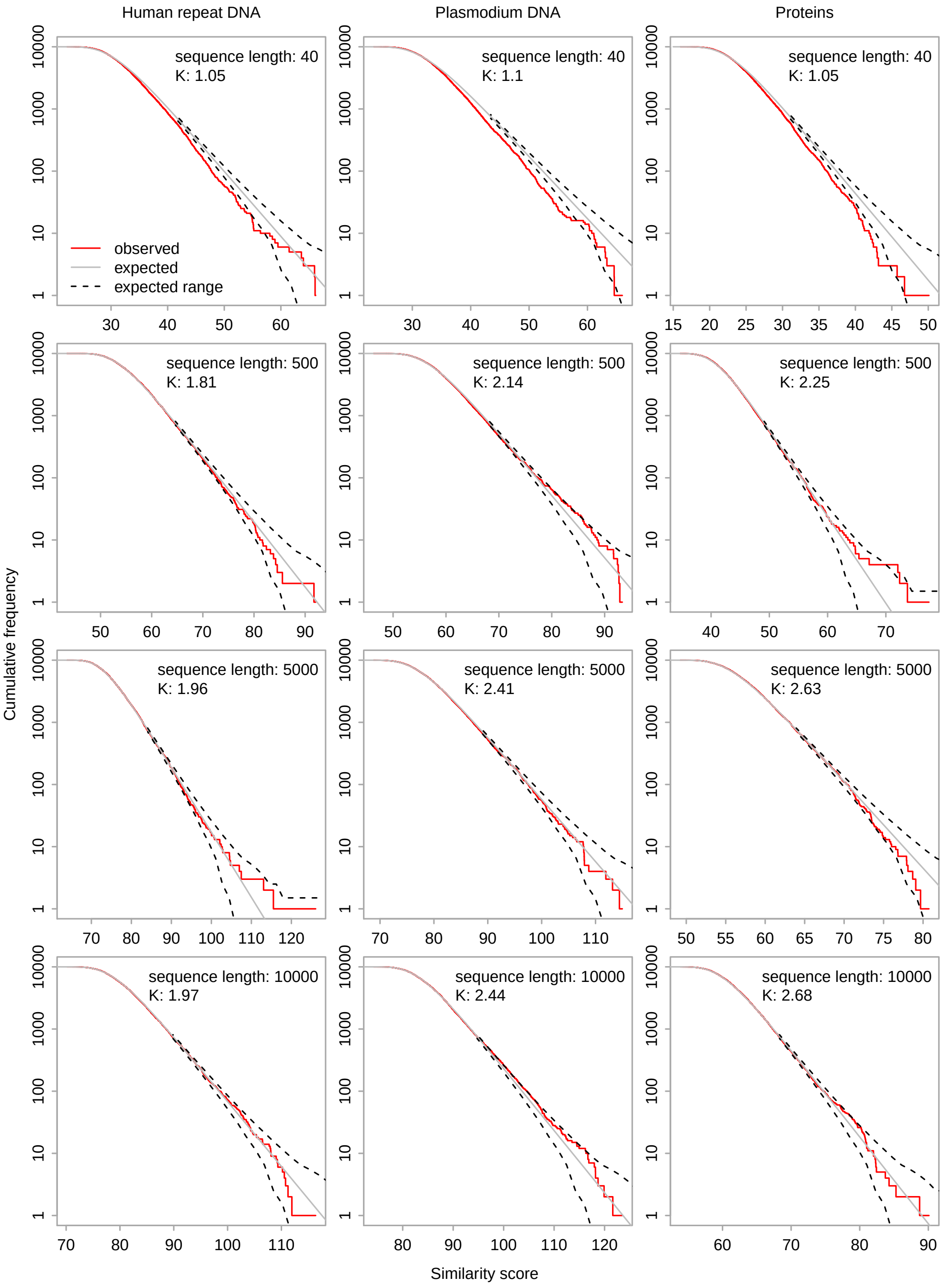


**Figure S1:** Non-heuristic similarity score ($s_{\max}$) between 10000 pairs of random i.i.d. sequences, for three sets of alignment parameters. The frequency has a 5% chance of being outside the dashed lines (2.5% each).

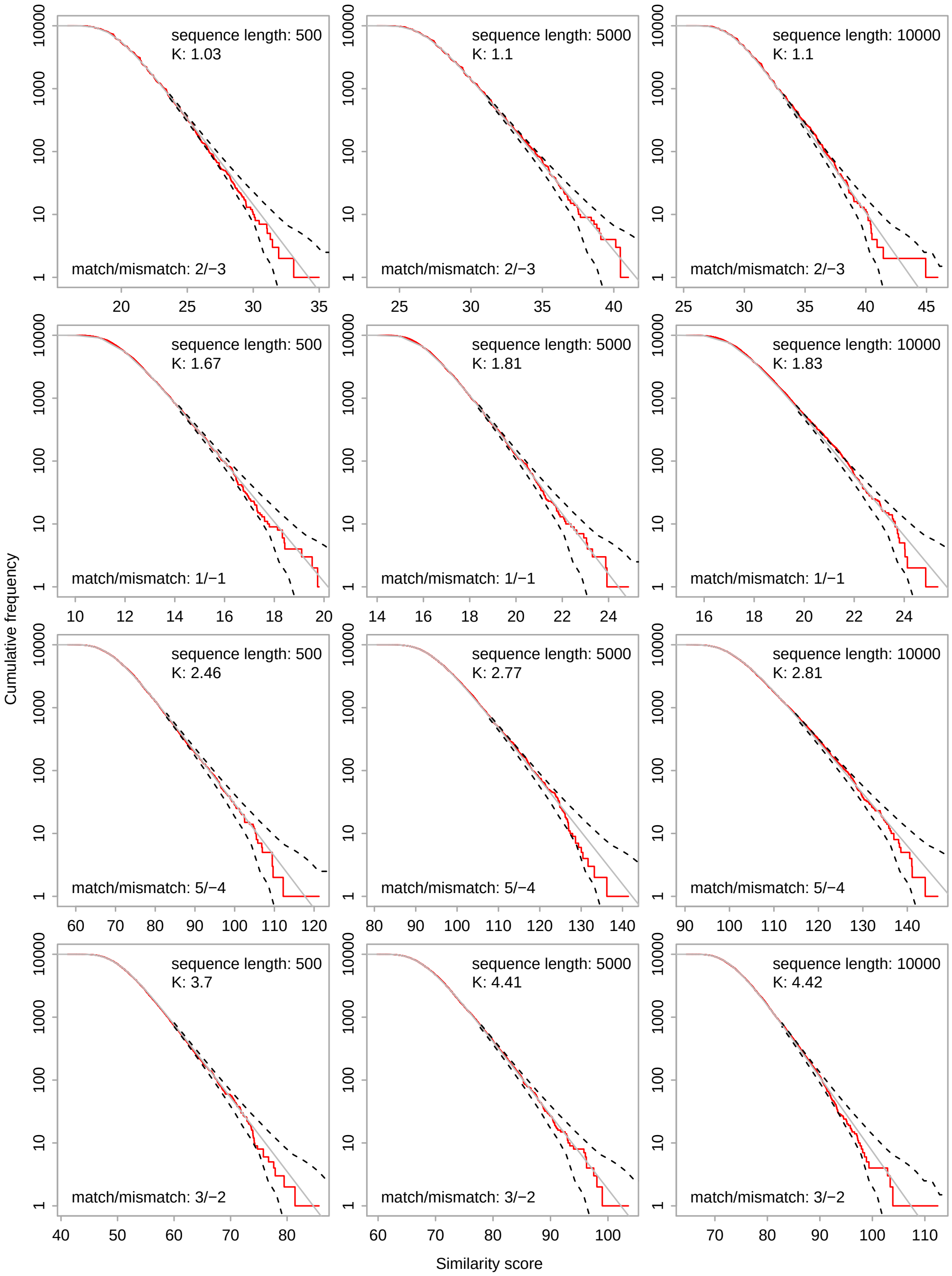


**Figure S2:** Non-heuristic similarity score ($s_{\max}$) between 10000 pairs of random i.i.d. sequences, for four sets of match/mismatch parameters. The frequency has a 5% chance of being outside the dashed lines (2.5% each).

4. Yao, Y & Frith, MC. Improved DNA-Versus-Protein Homology Search for Protein Fossils. *IEEE/ACM Transactions on Computational Biology and Bioinformatics* **20,** 1691–1699 (2023).
5. Frith, MC. How sequence alignment scores correspond to probability models. *Bioinformatics* **36,** 408–415 (2020).
6. Hamada, M, Wijaya, E, Frith, MC & Asai, K. Probabilistic alignments with quality scores: an application to short-read mapping toward accurate SNP/indel detection. *Bioinformatics* **27,** 3085–3092 (2011).
7. Frith, MC & Noé, L. Improved search heuristics find 20 000 new alignments between human and mouse genomes. *Nucleic Acids Research* **42,** e59–e59 (2014).
8. Frith, MC. A new repeat-masking method enables specific detection of homologous sequences. *Nucleic Acids Research* **39,** e23–e23 (2011).
9. Altschul, SF. A protein alignment scoring system sensitive at all evolutionary distances. *Journal of Molecular Evolution* **36,** 290–300 (1993).
10. Yu, YK & Hwa, T. Statistical significance of probabilistic sequence alignment and related local hidden Markov models. *Journal of Computational Biology* **8,** 249–282 (2001).
